# Mesoaccumbal glutamate neurons drive reward via glutamate release, but aversion via dopamine co-release

**DOI:** 10.1101/2022.03.17.484767

**Authors:** Shelley M. Warlow, Vivien Zell, Avery C. Hunker, Larry S. Zweifel, Thomas S. Hnasko

## Abstract

Ventral tegmental area (VTA) projections to the nucleus accumbens medial shell (NAc) drive reward-related motivation. Although dopamine neurons are predominant, a substantial glutamatergic projection is also present, and a subset of these populations can release both dopamine and glutamate. Optogenetic stimulation of VTA glutamate neurons supports self-stimulation, but can also induce place avoidance, even in the same assay. Here, we parsed the selective contribution of glutamate or dopamine co-release from VTA glutamate neurons to reinforcement and avoidance. We expressed Channelrhodopsin (ChR2) in VTA glutamate neurons, in combination with CRISPR/Cas9 to disrupt either the gene encoding vesicular glutamate transporter 2 (VGLUT2) or *Tyrosine hydroxylase* (*Th*). Selective disruption of VGLUT2 abolished optogenetic self-stimulation, but left real-time place avoidance intact, while CRISPR/Cas9 deletion of *Th* preserved optogenetic self-stimulation but abolished place avoidance. Our results demonstrate that glutamate release from VTA glutamate neurons is positively reinforcing, but that dopamine release from these same neurons can induce avoidance behavior.

## Introduction

As a principal region within mesocorticolimbic circuitry, the ventral tegmental area (VTA) significantly regulates reward-related motivation, aversion, and cognition. Although primarily composed of dopamine neurons, the VTA is a heterogenous structure containing GABA and glutamate neurons, as well as neurons containing multiple neurotransmitters (Hnasko et al., 2012; Pupe and Wallén-Mackenzie, 2015; Trudeau et al., 2014). Indeed, a subset of VTA dopamine neurons projecting to the medial nucleus accumbens (NAc) shell also express the type 2 vesicular glutamate transporter (VGLUT2) and release glutamate at NAc terminals (Kawano et al., 2006; Stuber et al., 2010; Tecuapetla et al., 2010).

Optogenetic stimulation of VGLUT2-expressing VTA neurons and their projections to medial NAc shell, lateral habenula, and ventral pallidum can promote positive reinforcement (Wang et al., 2015; Yoo et al., 2016). In self-stimulation assays, mice will perform instrumental actions such as nose poking or lever pressing to receive optogenetic stimulation of VTA glutamate neurons, suggesting increases in their activity is rewarding. VTA VGLUT2-mediated reinforcement persists despite manipulations that abolish concomitant dopamine release (Zell et al., 2020), suggesting dopamine release is not necessary for positive reinforcement mediated by VTA glutamate neurons. However, it has remained unknown whether glutamate release by these neurons is required for reinforcement behaviors.

VTA glutamate neurons have also been implicated in mediating aversive motivation, and it has been difficult to reconcile how mice will perform an instrumental action for optogenetic stimulation of VTA glutamate neurons in one assay, while simultaneously showing avoidance behavior in a different assay. For example, mice will avoid an arena paired with optogenetic stimulation of VTA glutamate neurons in a real time place procedure assay (Bimpisidis et al., 2019; Yoo et al., 2016; Zell et al., 2020). Consistent with this, mice will work to turn off stimulation in a negative reinforcement assay (Qi et al., 2016). We have previously suggested that the avoidance behavior may be a secondary consequence of the subjects’ preference for brief trains of optogenetic stimulation (<5s), and is a feature which distinguishes reinforcement responses to stimulation of VTA glutamate compared to VTA dopamine neurons (Yoo et al., 2016). However, another possibility is that glutamate and dopamine released from VTA glutamate projections to NAc shell mediate opposing effects. Indeed, increasing evidence suggests that dopamine release in NAc shell sub-regions relates to aversion (Badrinarayan et al., 2012; de Jong et al., 2019; Wenzel et al., 2015).

In the present study we tested how glutamate or dopamine release from VTA glutamate terminals in NAc contributes to instrumental reinforcement and place avoidance behaviors evoked by optogenetic stimulation. We show that selective disruption of glutamate release from VTA glutamate neurons via CRISPR-Cas9 abolished optogenetic self-stimulation, but that place avoidance behavior persisted. Conversely, disruption of dopamine release from VTA glutamate neurons abolished optogenetic-evoked place avoidance, while self-stimulation remained intact. Finally, disruption of both glutamate and dopamine-release from VTA glutamate projections abolished both positive reinforcement and avoidance behaviors. Our results suggest that mesoaccumbens glutamate release is a potent reinforcer, but that dopamine co-release from the same cells is, surprisingly, aversive.

## Results

### CRISPR/Cas9 disruption of glutamate transmission from mesoaccumbens projections

To disrupt glutamate release from VTA neurons, we used a single adeno-associated virus (AAV) vector for Cre recombinase-dependent expression of *Staphylococcus aureus* Cas9 (SaCas9) plus a single guide RNA (sgRNA) targeted to the gene encoding type 2 vesicular glutamate transporter (VGLUT2/*Slc17a6*) (Hunker et al., 2020). When injected into VTA of VGLUT2-Cre mice, Cas9 is expressed selectively in glutamate neurons to generate indel mutations resulting in loss of VGLUT2. To quantify the extent to which VGLUT2 function was lost, we co-injected AAV to achieve Cre-dependent Channelrhodopsin (ChR2) expression, in combination with an SaCas9 AAV targeted to VGLUT2 (sgVGLUT2) or a control targeted to the Rosa26 locus that is without functional consequence (sgControl) (***Figure 1a-b***). After 6 weeks, we recorded optogenetic-evoked excitatory postsynaptic currents (oEPSCs) from medium spiny neurons in NAc medial shell, where VTA glutamate neurons send dense projections. Cas9 disruption of VTA VGLUT2 significantly reduced the amplitude of oEPSCs compared to sgControl mice; oEPSCs were 68±9 pA in sgControl mice whereas sgVGLUT2 oEPSCs were 9±1 pA (t_26_=6.2, p<.0001), both of which were DNQX-sensitive (F_1,10_=42.0, p<.0001) (***Figure 1c-d***), indicating that CRISPR/Cas9 mediated disruption of VGLUT2 dramatically reduced glutamate release from VTA terminals in medial NAc shell.

**Figure 1.**
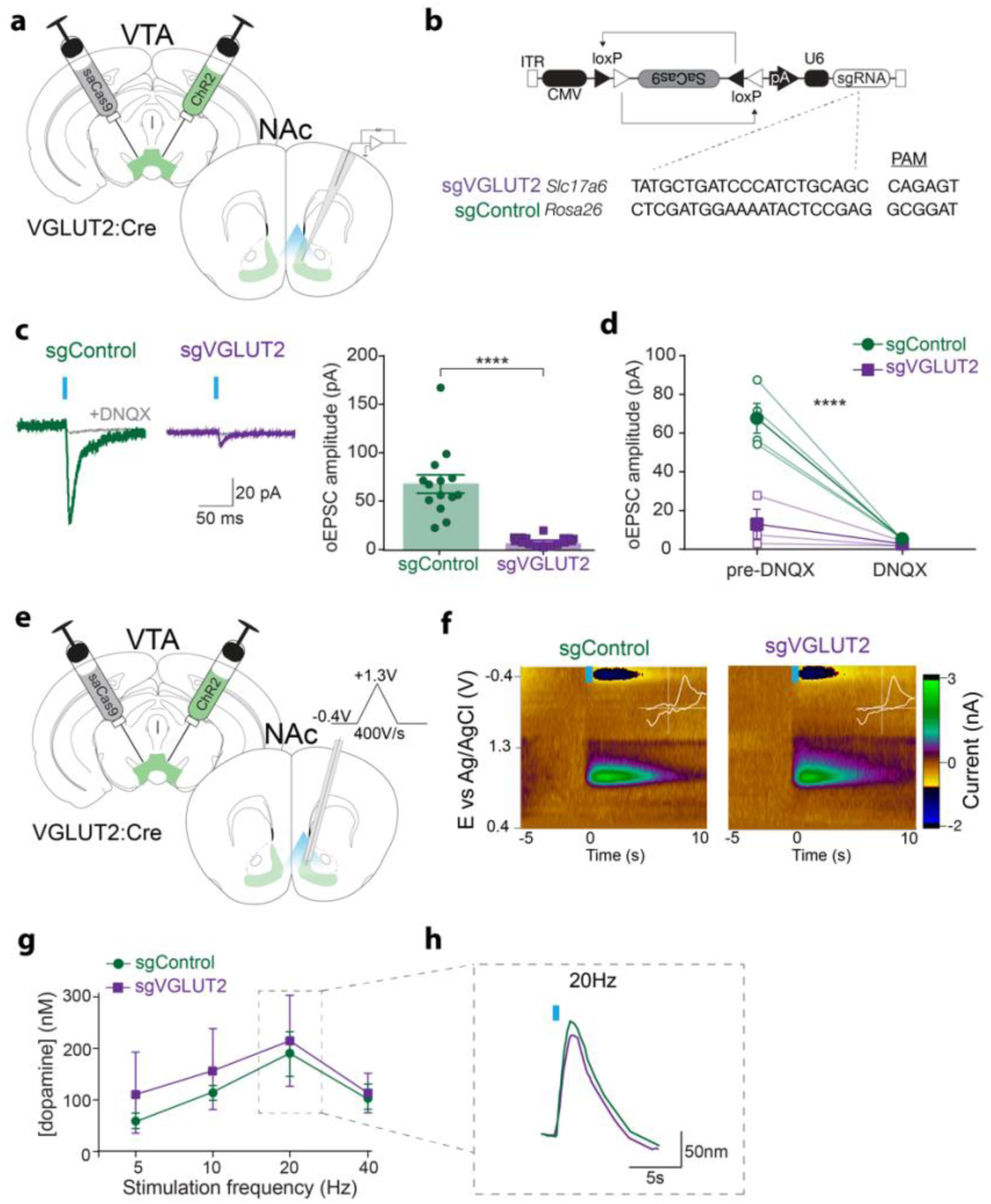
CRISPR-Cas9 deletion of VGLUT2 in VTA glutamate neurons abolishes VTA → NAc glutamate release but maintains dopamine co-release. a) Illustration of viral injections into VTA and of recording patch pipette in post synaptic NAc neurons. b) Design of AAV1-FLEX-SaCas9-U6-sgVGLUT2 and AAV1-FLEX-SaCas9-U6-sgROSA26, with guide sequences for each plus their associated PAM sites. c) Example traces of excitatory postsynaptic currents evoked by a 5-ms pulse of optogenetic stimulation (oEPSC) at V_h_=-60mV. oEPSC amplitudes were abolished in cells from sgVGLUT2 mice (n=14 sgVGLUT2, n=14 sgControl; unpaired t-test: p<0.0001). d) oEPSCs were blocked by bath-application of DNQX in both sgControl and sgVGLUT2 groups (n=7; main effect of treatment: p<0.0001). e) Illustrations of viral injections into VTA and carbon fiber electrode recordings in medial NAc shell applying a triangular waveform from −0.4 to 1.3V at a rate of 400V/s. f) Example color plots of phasic dopamine transients in response to optogenetic stimulation (20Hz, 1s). White inset depicts cyclic voltammogram (x-axis: −0.4 to 1.3 V, y-axis: −2 to 3 nA). g) Peak concentration of dopamine in response to increasing stimulation frequencies (n=4 sgVGLUT2 mice, n=4 sgControl mice). h) Example traces of dopamine concentration in response to 20-Hz optogenetic stimulation. ****p<0.0001.

Approximately 15-20% of VTA glutamate neurons co-express the dopamine marker *Tyrosine hydroxylase* (*Th*) and thus have the capacity to co-release dopamine in the NAc medial shell (Alsiö et al., 2011; Kawano et al., 2006; Stuber et al., 2010; Yamaguchi et al., 2011; Zell et al., 2020). To verify selective disruption of glutamate release while leaving dopamine co-release intact, we measured optogenetic-evoked dopamine in NAc medial shell using fast scan cyclic voltammetry (***Figure 1e***). Train stimulation (473nm, 1-s duration, 5-40 Hz) evoked dopamine release from VTA glutamate terminals in NAc medial shell, with a maximal effect around 20 Hz (main effect of frequency: F_3,18_=13.5, p<.0001), and no significant differences in peak dopamine were detected between sgControl and sgVGLUT2 conditions (main effect of group: F_1,6_=0.23, p=0.65) (***Figure 1f-h***). These results establish that Cas9 disruption of VGLUT2 in VTA glutamate neurons selectively abolished glutamate release without disrupting concomitant dopamine co-release in NAc.

### Mesoaccumbens glutamate release mediates positive reinforcement

Previous work demonstrated that mice nose poke to earn optogenetic stimulation of VTA glutamate neurons including cell bodies in VTA as well as their projections to NAc, lateral habenula, or ventral pallidum (Wang et al., 2015; Yoo et al., 2016; Zell et al., 2020). However, there is no direct evidence that it is the glutamate signal that is responsible for this behavioral reinforcement, and one report suggests that mesoaccumbens glutamate release activates NAc interneurons to promote aversion (Qi et al., 2016). To assess this, we used two separate reinforcement assays, a nose-poke and a place-based assay. Cre-dependent ChR2 was expressed in VTA of VGLUT2-Cre mice in combination with either sgVGLUT2 or sgControl. A third group received sgControl plus a control for the opsin (YFP), and in all groups optic fibers were placed to activate VTA terminals in NAc (***Figure 2a-b; Supplemental Figure 1a-b***).

**Figure 2.**
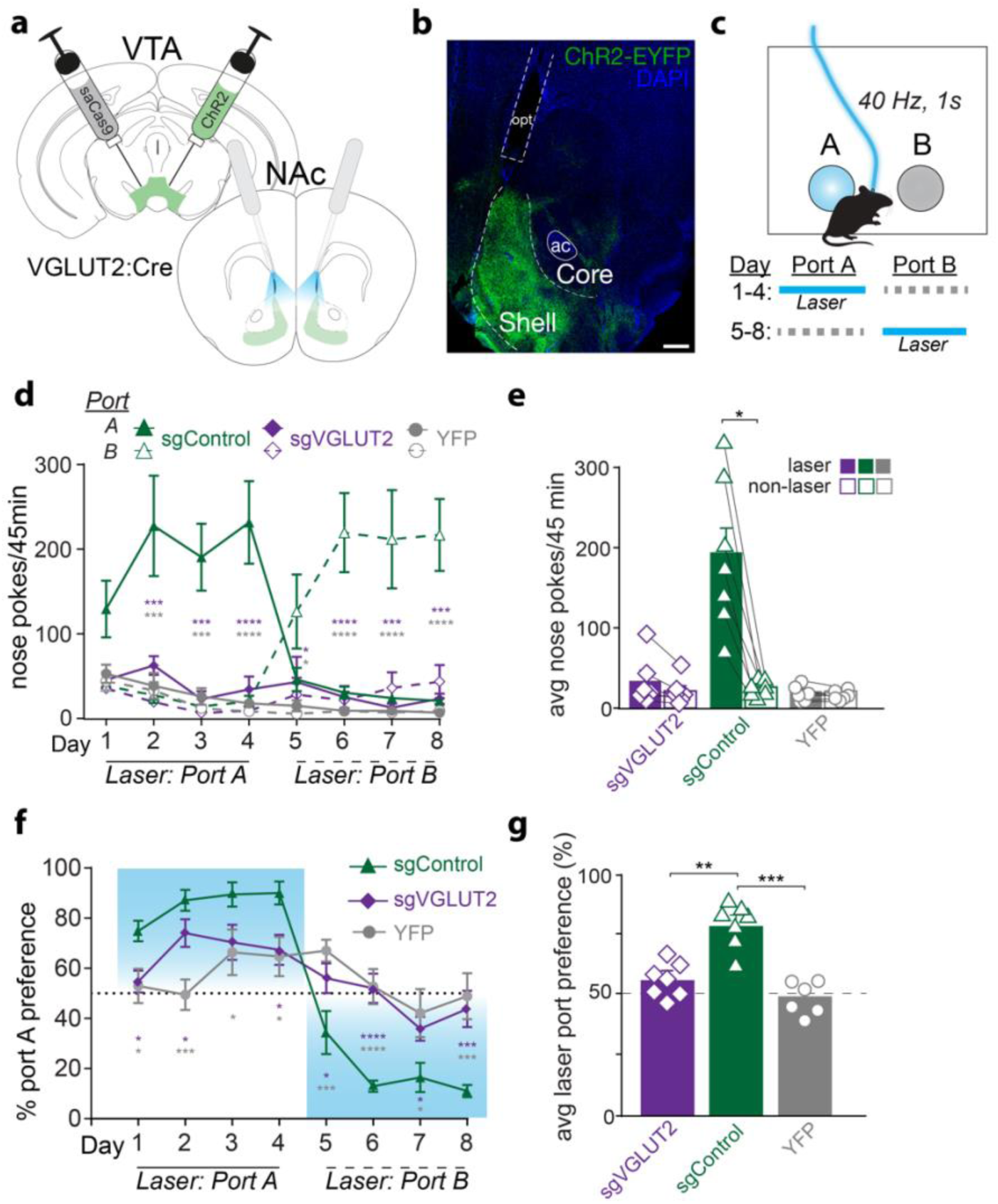
VTA glutamate projections require glutamate release to promote intracranial self-stimulation. a) Illustration of injection into VTA and optogenetic stimulation of VTA terminals in NAc with bilateral fiber implants. b) Image of coronal section showing ChR2 (ChR2-EYFP, green) and DAPI (blue) expression in NAc. Scale bar=200μm, anterior commissure (ac), optic fiber track (opt). c) Schematic of 2-nosepoke intracranial self-stimulation assay. Active nose pokes into laser-paired port earned 40 Hz, 1 s, 10 mW blue light. Location of laser-paired port was reversed starting Day 5. d) Active nose pokes at port A and B during each 45 min session. Sidak’s multiple comparisons tests comparing laser-paired pokes between sgControl (n=7) and sgVGLUT2 (n=7; purple asterisks) or between sgControl and YFP (n=6; grey asterisks). e) Laser and non-laser nosepokes averaged across all training days. Average laser nose pokes were greater than non-laser nosepokes only among sgControl mice (n=7; Sidak multiple comparisons test, t=7.1, p<0.05). f) Percent preference for Port A during each training session. Holm-Sidak’s multiple comparisons tests comparing laser-paired pokes between sgControl (n=7) and sgVGLUT2 (n=7; purple asterisks) or between sgControl and YFP (n=6; grey asterisks). g) Percent laser port preference averaged across all training days. Average laser port preference among sgControl mice (n=7) was greater than sgVGLUT (n=7; Tukey’s multiple comparison test, q=7.6, p<0.001) and YFP (n=6; Tukey’s multiple comparison test, q=9.5, p<0.0001) groups. *p<0.05, **p<0.01, ***p<0.001, ****p<0.0001

In an intracranial self-stimulation (ICSS) nose-poke assay, each nose poke into the active port (at fixed ratio 1) earned laser stimulation (40 Hz, 5-ms pulse width, 1-s duration, 473nm) plus a 1-s auditory tone. Nose pokes into an inactive port during the same session also triggered the auditory tone, but no laser (***Figure 2c***). ChR2 mice with intact mesoaccumbens glutamate release (sgControl) self-stimulated, making greater active vs. inactive nose pokes (main effect of port: F_1,6_=15.1, p=.008; ***Figure 2d***). Starting on day 5 the location of the active and inactive ports were reversed for the remainder of testing and sgControl mice tracked the laser location from the first day, making significantly more nose pokes for the new active port than inactive port (main effect of port: F_1,6_=11.4, p=.009). By contrast, sgVGLUT2 mice did not self-stimulate laser, such that that active nose pokes by sgControl mice were significantly greater compared to both YFP-expressing mice (without ChR2) and sgVGLUT2 mice across all days (main effect of group: F_2,17_=12.7, p=0.0004; ***Figure 2e***). Correspondingly, sgControl mice displayed an ∼80% preference for the active vs. inactive port which was significantly greater than both sgVGLUT2 and YFP mice (main effect of group: F_2,17_=12.2; p=0.0005), both of which displayed no preference for active vs. inactive ports (***Figure 2f-g***). These results establish that activation of NAc-projecting VTA glutamate neurons supports positive reinforcement in a manner that depends on VGLUT2-mediated glutamate release.

### Mesoaccumbens glutamate release does not mediate aversion

Previous work has shown that using a real-time place procedure (RTPP) with delivery of laser stimulation in one of two compartments, mice spent less time in a compartment paired with activation of mesoaccumbens glutamate neurons (Bimpisidis et al., 2019; Qi et al., 2016; Yoo et al., 2016; Zell et al., 2020). However, mice simultaneously showed an increase in approach rate, or entries, into the laser-paired compartment, and heat maps depicted a majority of time spent in the middle chamber (Yoo et al., 2016; Zell et al., 2020). These data suggest that activation of VTA glutamate neurons may provoke both rewarding and aversive responses.

Thus, we next used RTPP to assess whether glutamate release by VTA glutamate neurons is necessary for this distinct signature of positive reinforcement that includes apparent place avoidance but with increased approach rate. After a baseline day without laser delivery, mice were allowed to trigger laser stimulation by entering one of two compartments (***Figure 3a***). On day 4, the laser-paired side was switched for the remainder of test days. While YFP-expressing control mice spent similar amounts of time in each compartment, both sgVGLUT2 and sgControl mice spent less time in the laser-paired compartment (side x group interaction: F_2,17_=3.88, p=.04). These data indicate that place avoidance evoked by optogenetic stimulation of mesoaccumbens glutamate neurons does not depend on VGLUT2-mediated glutamate release (***Figure 3b-c***).

**Figure 3.**
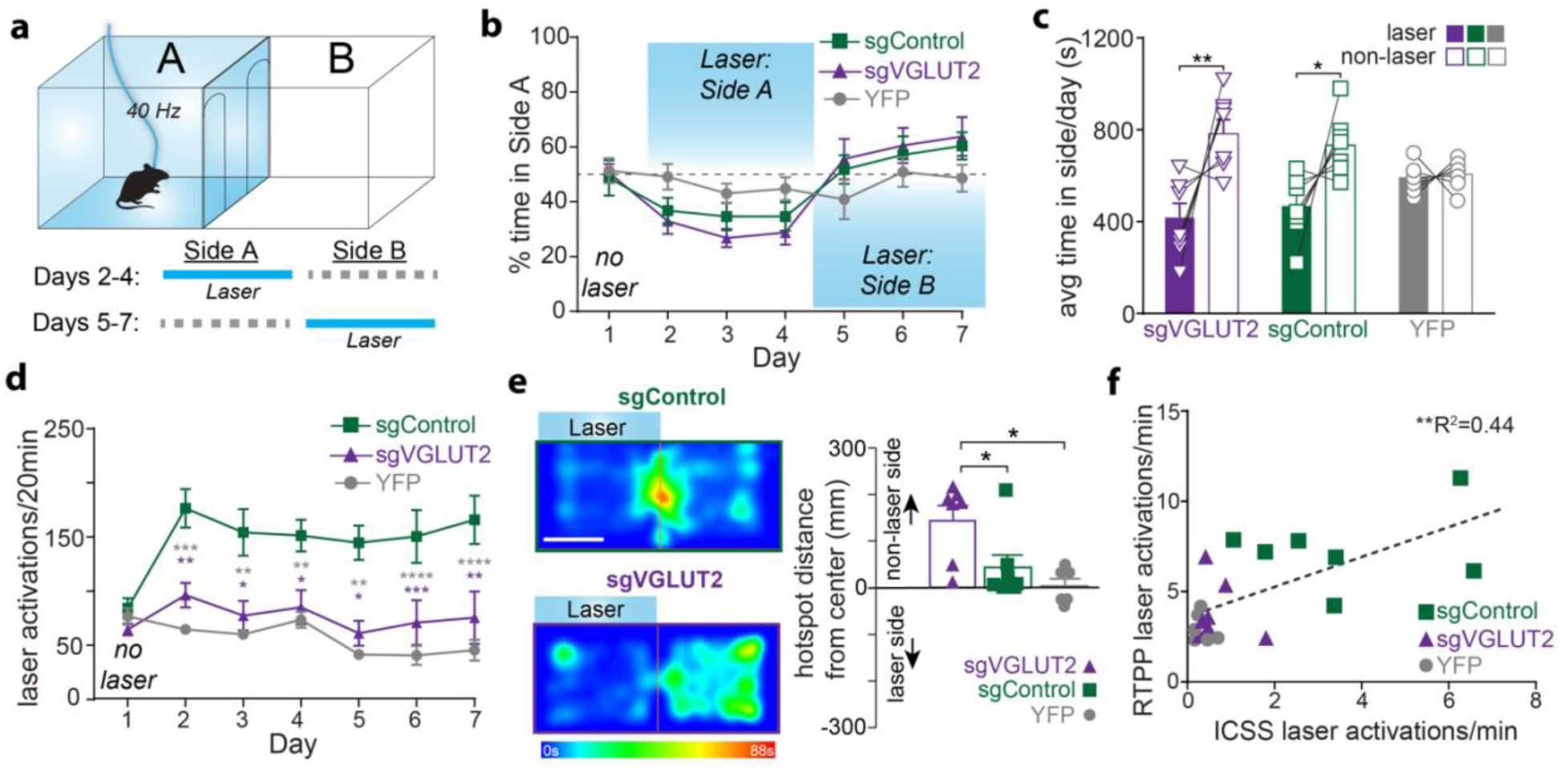
VTA glutamate projections require glutamate release to promote laser activations but not place avoidance. a) Schematic of real time place procedure (RTPP). Starting on Day 2, entries into laser-paired port earned 40 Hz, 10 mW blue light. Location of laser-paired side was reversed starting Day 5. b) Percent time in Side A during each 20-min daily session. c) Time spent in laser and non-laser sides averaged across all laser sessions (Days 2-7). Average time spent in laser side was lower than time in non-laser side among both sgControl mice (n=7; Holm-Sidak’s multiple comparisons test, p<0.05) and sgVGLUT2 mice (n=7; Holm-Sidak’s multiple comparisons test, p<0.01), but not among YFP mice (n=6; Holm-Sidak’s multiple comparisons test, p>0.05). d) Laser activations (identical to number of entries into laser-paired compartment) during each daily 20-min session. Baseline session on Day 1 shows entries into compartment but no laser was delivered. Tukey’s multiple comparisons tests comparing laser activations between sgControl (n=7) and sgVGLUT2 (n=7; purple asterisks) or between sgControl and YFP (n=6; grey asterisks). e) Example heat maps from Day 4 of RTPP. White scale bar=135 mm. Bar graph depicts distance from center for each mouse’s hotspot (most time spent) on day 4. Hotspots of sgVGLUT2 (n=7) were further away from the center of the compartments than sgControls (n=7; Tukey’s multiple comparisons test, p<0.05) and YFPs (n=6; Tukey’s multiple comparisons test, p<0.05). f) Average laser activations earned per minute across ICSS 2-nosepoke sessions plotted against average laser activations earned per minute across RTPP sessions (n=20; Pearson correlation, r=0.66, p<0.002).

While the place avoidance was similar between sgVGLUT2 and sgControl mice, sgVGLUT2 mice failed to show the signature increase in approach rate in this assay. Indeed, we found that, the number of laser activations (caused by entry into the laser-paired compartment) increased across sessions among the sgControl mice but did not increase among sgVGLUT2, which were similar to YFP levels (day x virus interaction: F_12,102_=4.59, p<.0001; ***Figure 3d***). Furthermore, heat maps depict sgControl mice spent a majority of time near the center of both compartments, presumably as a consequence of their making frequent brief entries to the laser-paired chamber, whereas sgVGLUT2 subjects spent most time in the non-laser-paired compartment further from the center line (distance from center, F_2,17_=6.03, p<.01; ***Figure 3e***). The failure of sgVGLUT2 mice to show the signature increase in laser compartment entries (and subsequent laser activations) is unlikely to reflect a baseline change in locomotion because distance traveled in an open field chamber was similar between the groups (t_12_=0.46, p=0.65) (***Supplemental Figure 2a***). Instead, the increase in chamber entries appears to reflect mice seeking short bouts of laser reinforcement. Consistent with this hypothesis, the number of laser activations mice made in the RTPP is well correlated with the number of laser activations in the nose poke ICSS assay (Pearson r=0.80, p<0.0001) (***Figure 3f***).

Disruption of VTA glutamate release selectively abolished laser self-stimulations in the nose poke ICSS and RTPP assays, suggesting that glutamate release from VTA glutamate neurons mediates positive reinforcement. The lack of positive reinforcement was unlikely due to a generalized deficit in basic reward learning. When tested in an instrumental lever pressing task to earn food pellets (FR1), both sgVGLUT2 and sgControl mice demonstrated similar levels of lever pressing to earn a pellet (main effect of group: F_2,17_=0.81, p=0.46) (***Supplemental Figure 2b***). Thus, glutamate release from VTA VGLUT2 neurons is responsible for optogenetic reinforcement when these neurons are targeted, but loss of VGLUT2 from these cells is not essential to learn a similar instrumental task for food reinforcers.

### Dopamine co-release from mesoaccumbens glutamate neurons mediates avoidance

Since place avoidance in the RTPP did not depend upon glutamate release from VTA glutamate neurons, we next tested the effects of disrupting dopamine co-release from VTA glutamate neurons on self-stimulation and place avoidance. We expressed Cre-dependent ChR2 in combination with Cre-dependent SaCas9 i) targeted to *Th* to disrupt dopamine co-release (sgTH), ii) targeting both *Th* and VGLUT2 to abolish both glutamate and dopamine release (sgTH+sgVGLUT2), iii) targeting the Rosa26 control (sgControl); and iv) a fourth group that expressed YFP instead of ChR2. (***Figure 4a***).

**Figure 4.**
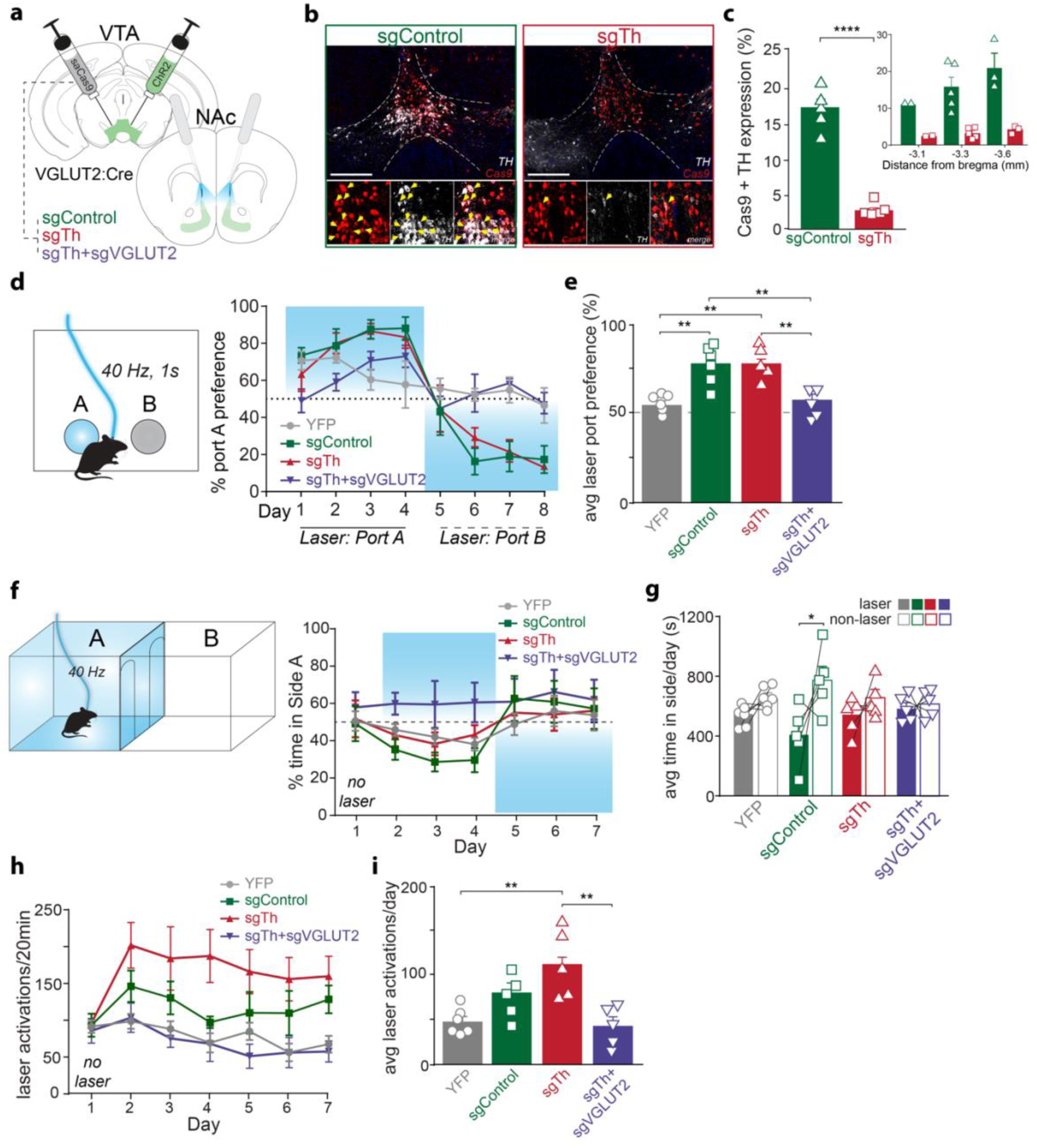
VTA glutamate projections require dopamine release to promote place avoidance but not self-stimulation. a) Illustration of injections in VTA and optic fiber placements above NAc. b) Images of coronal sections showing Cas9 (red) and TH (white) expression in VTA of sgControl and sgTh mice. Scale bar=300 μm. Yellow arrows indicate cells co-expressing Cas9 and Th. c) Percent of Cas9 cells in VTA that also express Th were higher among sgControls (n=5 mice) than sgTh (n=5 mice; unpaired t-test, t_8_=9.6, p<0.0001). Right inset graph depicts percent Cas9 and Th co-expression across anterior-posterior VTA sites. d) Schematic of 2-nosepoke ICSS assay. Percent preference for Port A across daily training sessions. Location of laser port was reversed starting on day 5. e) Holm-Sidak’s multiple comparison tests show average laser port preference among sgControl mice (n=5) was greater than YFP (n=6; t=4.3, p<0.01) and sgTh+sgVGLUT2 mice (n=5; t=4.0, p<0.01), but similar to sgTh (n=5; t=0.59, p>0.05), while sgTh mice also showed greater laser preference than YFP (t=3.7, p<0.01) and sgTh+sgVGLUT2 mice (t=3.4, p<0.01). f) Average laser nose pokes were greater than non-laser nosepokes among sgControl mice (n=5; Sidak multiple comparisons test, p=0.0002) and sgTh mice (n=5; Sidak multiple comparisons test, p=0.048). g) Schematic of RTPP assay. Percent time in Side A during each 20-min daily session. No laser was delivered on day 1, and location of laser side was reversed on day 5. h) Average time spent in laser side was lower than time in non-laser side among sgControl mice (n=5; Holm-Sidak’s multiple comparisons test, t=2.9, p<0.05) but not among sgTh mice (n=5; t=1.1, p>0.05), sgTH+sgVGLUT2 mice (n=5; t=0.31, p>0.05), or YFP mice (n=5; t=1.2, p>0.05). i) Laser activations earned during each daily 20-min session. j) Laser activations averaged across laser sessions in sgTh mice (n=5) were higher than YFP mice (n=6; Tukey’s multiple comparisons test, t=3.7, p<0.01) and sgTh+sgVGLUT2 mice (n=5; Tukey’s multiple comparisons test, t=3.8, p<0.01).

To validate the *Th* disruption by CRISPR/Cas9, we quantified co-expression of Cas9-expressing cells with those expressing TH (***Figure 4b***). Immunohistochemical analysis revealed that 17.2±1.4% of VGLUT2-Cre neurons expressing Cas9 in VTA also co-expressed TH in sgControl mice, whereas only 3.1±0.5% co-expressed TH among sgTh mice (t_8_=9.63, p<0.0001) (***Figure 4c***), demonstrating efficient disruption of TH expression in VTA glutamate neurons.

To test the effect of dopamine release from VTA glutamate neurons on positive reinforcement, we first tested mice in the ICSS nose-poke assay (***Figure 4d***). Both sgControl and sgTh mice self-stimulated laser, making greater number of active vs. inactive nose pokes. In contrast, both YFP and sgTh+sgVGLUT2 groups made fewer active nose poke responses and showed no preference for active vs. inactive (group x port interaction: F_3,18_=4.46, p=0.017). As a result, sgControl and sgTh mice displayed a high preference for the laser-paired port across training days compared to equal preference among YFP and sgTH+sgVGLUT2 mice (main effect of group: F_3,18_=10.0, p=0.0004) (***Figure 4e***). These results indicate that self-stimulation of VTA glutamate projections to NAc occur despite abolishing dopamine co-release, consistent with our previous findings (Zell et al., 2020), and further show that positive reinforcement is abolished when both glutamate and dopamine release are abolished.

We next tested the contribution of dopamine release by VTA glutamate neurons to place avoidance using the RTPP task (***Figure 4f***). sgControl mice avoided the laser-paired compartment, spending less time in the laser-paired side than non-laser side across training days, and switching preference when the active side was switched on day 5. By contrast, both sgTh and sgTh+sgVGLUT2 mice showed no place avoidance or preference, spending equal time in both compartments, without switching preference when the active side was switched (group x side interaction: F_3,18_=9.1, p=0.007) (***Figure 4g***). However, while sgTh mice showed no place avoidance, they continued to show an increased number of entries into the laser side, earning an equivalent number of increased laser activations per session as sgControls (***Figure 4h-i***). In contrast, the number of laser activations by sgTh+sgVGLUT2 were comparable to YFP opsin controls. We observed similar effects of sgTH-mediated dopamine disruption in a separate cohort of mice (***Supplemental Figure 3***). Together, these results suggest that dopamine co-release from mesoaccumbens glutamate neurons mediates an aversive signal that leads to place avoidance, while glutamate release from these cells promotes positive reinforcement.

## Discussion

Stimulation of VTA glutamate neuron projections to NAc medial shell simultaneously promotes positive reinforcement but also place avoidance. Here, we demonstrate that glutamate release by VTA VGLUT2 neurons is necessary for stimulation-evoked positive reinforcement, while dopamine release from the same population is required for avoidance. Selectively disrupting VGLUT2 from VTA neurons through CRISPR-Cas9 endonucleases abolished presynaptic glutamate release without disruption of co-released dopamine. As a result, self-stimulation in both nose-poke and place-based (laser-side entries) assays was abolished. Yet VGLUT2 disruption did not alter place avoidance caused by VTA VGLUT2 stimulation. By contrast, selective disruption of dopamine co-release from VTA glutamate neurons left positive reinforcement intact but abolished place avoidance, indicating that dopamine released from VTA glutamate neurons promotes aversion. We further demonstrate that CRISPR-Cas9 deletion of both TH and VGLUT2 completely abolishes both self-stimulation and avoidance behaviors. These results demonstrate that glutamate and dopamine released by NAc-projecting VTA glutamate neurons differentially contribute to reward and aversion.

### Mesoaccumbal glutamate release promotes positive reinforcement

Optogenetic activation of excitatory inputs to the NAc from amygdala, hippocampus, and cortex have been repeatedly shown to reinforce behaviors (Britt et al., 2012; LeGates et al., 2018; Otis et al., 2017; Stuber et al., 2011). Our results are consistent with prior works and support a prominent role for excitatory inputs to NAc from VTA in positive reinforcement (Wang et al., 2015; Yoo et al., 2016; Zell et al., 2020). Our findings are also consistent with studies showing that conditional deletion of VGLUT2 from VTA dopamine neurons disrupts psychomotor sensitization and alters reward-seeking behaviors (Alsiö et al., 2011; Birgner et al., 2010; Hnasko et al., 2010; Papathanou et al., 2018), that photoinhibition of VTA glutamate neurons disrupts cue-reward associations (Yau et al., 2016), and that VTA glutamate neurons are activated by rewards and reward-predictive cues (McGovern et al., 2021; Root et al., 2020).

On the other hand, disrupting glutamate release from dopamine neurons does not blunt optogenetic self-stimulation of dopamine neurons (Wang et al., 2017). The most likely explanation for this is that only a minority of VTA dopamine neurons express VGLUT2, and that stimulation of the global population of VTA dopamine neurons is sufficient to support strong positive reinforcement independent of glutamate co-release from a minority subset of dopamine neurons. We have also shown that when comparing effects of stimulating the global pool of VTA glutamate versus the global pool of VTA dopamine neurons, mice prefer brief trains of VTA glutamate neuron stimulation (<5 s), but prefer more sustained trains of VTA dopamine neuron stimulation (>5 s) (Yoo et al., 2016). Thus, the release of glutamate from VTA neurons promotes reward-related motivation that is distinct and independent from dopamine.

### Positive reinforcement and avoidance are separable features of VTA glutamate neuron signaling

While stimulation of mesoaccumbal glutamate projections promotes self-stimulation and approach behavior, stimulation of these projections simultaneously results in place avoidance during real time place assays, and VTA glutamate neurons increase their intrinsic activity in response to both rewarding and aversive stimuli (Qi et al., 2016; Root et al., 2020; Yoo et al., 2016; Zell et al., 2020). Together, these data support a role for VTA glutamate neurons not only in reward, but also in aversion. Prior work suggested a possible role for glutamatergic activation of NAc parvalbumin-expressing (PV) interneurons in driving aversive behaviors (Qi et al., 2016). However, a more recent study showed that direct activation of NAc PV interneurons elicits a conditioned place preference while their inhibition produces conditioned avoidance (Chen et al., 2019), suggesting that avoidance is unlikely to be mediated via glutamatergic activation of PV interneurons. It is additionally unclear whether NAc PV neurons are a prominent post-synaptic target of VTA glutamate neurons, whose connections to both dopamine D1- and D2-receptor-containing medium spiny neurons and cholinergic interneurons in NAc have been better established (Chuhma et al., 2014; Mingote et al., 2015; Zell et al., 2020).

Our results propose an alternate explanation for the mixed positively and negatively reinforcing effects observed upon activation of VTA glutamate neurons, while also explaining why at least a subset of VTA glutamate neurons (those that co-release dopamine) may be activated by both rewarding and aversive stimuli. We propose that, like other glutamatergic inputs to NAc, glutamate released from VTA terminals in NAc is positively reinforcing. But that dopamine, which is co-released from a subset of these cells, leads to an aversive response. Neurotransmitter-defined aversion vs. reward may also occur in other targets of VTA glutamate projections, for example through the co-release of GABA and glutamate in ventral pallidum or lateral habenula (Lammel et al., 2015; Root et al., 2014; Yoo et al., 2016).

### VTA glutamate projections require dopamine co-release to promote avoidance

While dopamine is best known for supporting reward-related motivation, growing evidence points to diverse roles for dopamine across heterogeneous striatal sub-regions and cell-types (de Jong et al., 2022; Wenzel et al., 2015). Specifically, it has been repeatedly established that some dopamine neurons are activated in response to aversive stimuli (Brischoux et al., 2009; Matsumoto and Hikosaka, 2009; Ungless et al., 2004). In particular, the medial NAc shell, which receives the densest fraction of glutamatergic fibers from VTA (Hnasko et al., 2012; Taylor et al., 2014), is an apparent hotspot for dopamine release evoked by aversive stimuli. For example, multiple studies reported dopamine release in the medial NAc shell in response to an aversive stimulus (e.g., foot shock or threatening odor) or it’s associated cue; this in contrast to lateral NAc shell or core, where dopamine release decreased in response to aversive stimuli (Badrinarayan et al., 2012; de Jong et al., 2019). Mesolimbic dopamine is also critical for aversive processes such as innate defensive behaviors and fear conditioning (Fadok et al., 2009; Faure et al., 2008; Richard and Berridge, 2011; Zweifel et al., 2011), and one recent study showed a role for VTA glutamate neurons in mediating defensive responses to a looming threat (Barbano et al., 2020).

Our present findings demonstrate for the first time that mesolimbic dopamine, specifically co-released by VGLUT2-expressing VTA glutamate neurons, induces avoidance behavior elicited by activation of these neurons. Thus, the source of dopamine evoked in response to aversive stimuli is likely to include dopamine neurons that also release glutamate. Yet, whether dopamine release from non-glutamate dopamine neurons that project to medial NAc shell also contribute to aversive responses is unknown.

## Conclusion

Together, our data demonstrate that VTA glutamate projections promote positive reinforcement through release of glutamate, and simultaneously promote avoidance that instead relies on co-release of dopamine from these cells. This evidence further highlights mesolimbic contributions to reinforcement that are dopamine-independent, and expands our understanding of neurotransmitter-specific roles within co-releasing populations. Our findings also add to the growing evidence implicating opposing reward and aversive functions of mesolimbic dopamine signals and the importance of studying the role of dopamine and non-dopamine VTA sub-types. Indeed, the ability for a neuronal population to promote divergent functions at the level of the neurotransmitter has complex implications for understanding disorders involving dysregulated reward or aversion signaling, such as substance use or compulsive disorders.

## Methods

### Animals

Male and female mice were bred at University of California San Diego (UCSD) and group-housed on a 12-hour light/dark cycle, with food and water *ad libitum* unless otherwise noted. *Slc17a6*^*+/Cre*^ (VGLUT2cre) mice were initially obtained from Jackson Laboratory (Stock: 016963) and maintained back-crossed on to C57BL/6J. All experiments were in accordance with protocols approved by UCSD Institutional Animal Care and Use Committee.

### Viral Production

Production of AAV1 (AAV1-FLEX-ChR2-EYFP, AAV1-FLEX-EYFP, AAV1-FLEX-SaCas9-Ug-sgVglut2, AAV1-FLEX-SaCas9-U6-sgTh, and AAV1-FLEX-SaCas9-U6-sgRosa26) were as previously described (Gore et al., 2013). Briefly, pAAV shuttle plasmids were co-transfected with the packaging plasmid pDG1 (Grimm et al., 1998) into HEK293T/17 cells (ATCC) and viral particles were purified by cesium chloride gradient centrifugation. Viral particles were resuspended in Hank’s balanced saline solution and titers were calculated using gel electrophoresis and densitometry against a known standard.

### Stereotactic Surgeries

Mice > 6 weeks old were anesthetized with isoflurane (4% for induction; 1-2% maintenance) and placed in a stereotaxic frame (Kopf Instruments). AAV1-FLEX-ChR2-EYFP (8 × 10^12^ vg/mL) was combined with either AAV1-FLEX-SaCas9-U6-sgVglut2 (1.5 × 10^12^ vg/mL), AAV1-FLEX-SaCas9-U6-sgTh (1.8 × 10^12^ vg/mL), or AAV1-FLEX-SaCas9-U6-sgRosa26 (1.5 × 10^12^ vg/mL) such that ChR2-YFP constituted 1/7^th^ of the total volume and the respective SaCas9 constituted 6/7^th^ of the total volume. 400nL of this mixture was injected into the VTA of VGLUT2cre mice (Distance from Bregma in mm: −3.4 AP; +0.35 ML; −4.4 DV) at 100nl/min using a glass pipette attached to a microinjector (Nanoject 2, Drummond Scientific). Following viral infusion, the injector tip was kept in place for 10 min before slowly retracting. For optogenetic self-stimulation experiments, optic fibers (200μm core; Newdoon) were subsequently placed bilaterally above NAc medial shell at a 10° medio-lateral angle (+1.4 AP; ±1.13 ML; −3.81 DV). Optic fibers were secured with 4 skull screws and dental cement (Lang Dental Mfg). Mice were treated with topical antibiotic and with carprofen (5mg/kg; s.c.; Rimadyl) immediately following surgery and 24 hr later. Mice were allowed 6 weeks for recovery before experiments began.

### Immunohistochemistry

Mice were deeply anesthetized with sodium pentobarbital (200mg/kg; i.p.; VetOne) and transcardially perfused with 30-40 mL of phosphate-buffered saline (PBS) followed by 60-70 mL of 4% paraformaldehyde (PFA) at ∼7 mL/min. Brains were extracted and stored in 4% PFA overnight, followed by cryoprotection in 30% sucrose for 48-72 hr at 4°C. Brains were subsequently flash frozen in isopentane and stored at −80°C. 30-um sections were cut using a cryostat (Leica) and stored in PBS containing 0.01% sodium azide. For immunostaining, sections were blocked with 4% normal donkey serum (Jackson ImmunoResearch) in PBS containing 0.2% Triton-X 100 for 1 hour at room temperature. Sections were then incubated in sheep anti-TH (1:2000; Pelfreeze P60101-0) and rabbit anti-GFP (1:2000; Invitrogen A11122), or chicken anti-GFP (1:2000; Invitrogen A10262) and rabbit anti-HA (1:2000; Sigma H6908) overnight at 4°C. Following primary incubation, sections were rinsed in PBS 3 times for 10 min each and subsequently incubated in secondary antibodies conjugated to Alexa 488 (Donkey anti-rabbit; 1:400) and Alexa 647 (Donkey anti-sheep; 1:400), or Alexa 488 (Donkey anti-chicken; 1:400) and Alexa 594 (Donkey anti-rabbit; 1:400) (Jackson ImmunoResearch) for 2 hr at room temperature. Sections were then rinsed in PBS 3 times for 10 min each, mounted onto glass slides, and coverslipped with Fluoromount-G mounting medium (Southern Biotech) containing 0.5ug/mL DAPI (Sigma 268298).

Images were taken using a widefield epifluorescent microscope (Zeiss AxioObserver). Tiled images were taken at 10x magnification using appropriate filters and identical acquisition settings across all slides. Approximately 3-4 sections through rostral-caudal extent of VTA and 3-4 sections from NAc were imaged. Spread of ChR2 expression and optic fiber placements were mapped onto corresponding coronal sections in the Paxinos Mouse Brain Atlas using Adobe Illustrator (Version 23).

### Electrophysiological recordings in mouse brain slices

Adult mice 12-14 weeks old were deeply anesthetized with sodium pentobarbital (200mg/kg; i.p.; VetOne) and transcardially perfused with 15 mL ice-cold NMDG artificial cerebro-spinal fluid (aCSF) containing (in mM): 92 NMDG, 2.5 KCl, 1.25 NaH_2_PO_4_, 30 NaHCO_3_, 20 HEPES, 25 D-Glucose, 2 thiourea, 5 Na-ascorbate, 3 Na-pyruvate, 0.5 CaCl_2_, and 10 MgSO4, and continuously bubbled with carbogen (95% O_2_+ 5% CO_2_). Brains were then extracted and 200-μm coronal slices were cut using a vibratome (Leica) containing ice-cold NMDG-aCSF. Slices were then transferred to a recovery chamber containing NMDG-aCSF at 31°C for 20-30 min. A 2M Na^+^ spike-in solution (116mg/mL Na^+^ in NMDG-aCSF) was added to the recovery chamber in increasing volumes (from 250uL to 1mL) in 5 min increments for 25 min in order to achieve a controlled rate of reintroduction of Na^+^ into the chamber (Ting et al., 2018). 5 min after the last Na^+^-spiking solution, slices were then transferred into room temperature HEPES-aCSF containing (in mM): 92 NaCl, 2.5 KCl, 1.25 NaH_2_PO_4_, 30 NaHCO_3_, 20 HEPES, 25 D-Glucose, 2 thiourea, 5 Na-ascorbate, 3 Na-pyruvate, 2 CaCl_2_, 2 MgSO4, and continuously bubbled in carbogen. After >45 min recovery, slices were transferred to a recording chamber continuously perfused with carbogenated aCSF at a rate of 2-3 ml/min and maintained at 32°C by an in-line heater (Warren Instruments).

Patch-pipettes (4.5-6 MΩ) were pulled from borosilicate glass (Kings Precision Glass) using a gravity puller (Narishige). Pipettes were filled with a cesium-based internal solution containing (in mM): 130 D-Gluconic acid, 130 CsOH, 5 NaCl, 10 HEPES, 12 phosphocreatine, 3 Mg-ATP, 0.2 Na-GTP, 10 EGTA, at pH 7.25 and 285 mOsm. Epifluorescence was used to locate YFP-labelled VGLUT2 VTA terminals in NAc medial shell and subsequent visually guided patch recordings were made using infrared differential interference contrast (IR-DIC) illumination (A1 Examiner, Zeiss). A light-emitting diode (UHP-LED460, Prizmatix) under computer control was used to flash blue light through the light path of the microscope to activate ChR2. Recordings were made in whole cell voltage clamp using a Multiclamp 700B Amplifier (Axon Instruments), filtered at 2 kHz, and digitized at 10 kHz (Axon Digidata, Axon Instruments), and collected using pClamp v10 software (Molecular Devices).

Capacitance and series resistance were electronically corrected before recordings, and series resistance was monitored throughout recordings. Any cell in which series resistance changed >20% was discarded and excluded from analyses. To record excitatory postsynaptic currents (EPSCs), neurons were voltage clamped at −70 mV in whole cell configuration. A single 5-ms blue light pulse was applied every 45 s, and 10 light-evoked currents were averaged per neuron per condition. AMPA (α-amino-3-hydroxy-5-methyl-4-isoxazolepropionic acid) receptors were blocked using 6,7-dinitroquinoxaline-2,3-dione (DNQX, Sigma) dissolved in dimethyl sulfoxide and diluted by 1000 in aCSF for 10μM bath application.

### In vitro Fast-scan Cyclic Voltammetry Recordings

Coronal slices from adult mice were prepared as above with the exception that slices were cut to 300 μm thickness. Carbon fiber electrodes were prepared using 7 μM thick carbon fiber threaded through glass pipettes. Pipettes were pulled to seal around the carbon fiber, and subsequently cut so that ∼100μm of carbon fiber was exposed beyond the pipette. Pipettes were backfilled with 3M KCl, and a Ag/AgCl reference electrode was also placed in the recording chamber. The potential of the carbon fiber electrode was held at −4 V versus reference, ramped to 1.3 V, and back to −4 V at 400 V/s. This triangular waveform was first applied at 60 Hz for 10-15 min in the bath, and then at 10Hz for the duration of slice recordings. Dopamine transients were evoked optogenetically applied through the light path of the microscope as above, in 1-s trains at various frequencies (5-40 Hz). Optogenetic stimulations were applied every 2 min, in ascending order of frequency, with three repetitions for each frequency. Data were collected and analyzed using TarheelCV software. The amplitude of evoked dopamine transients were measured at the site of peak oxidation (0.6-0.7 V), and averaged across the 3 recordings for each frequency. To estimate dopamine concentration, each electrode was calibrated to 1000 nM dopamine (Alfa Aesar A11136) prepared fresh daily.

### 2 nose-poke optogenetic self-stimulation procedure

Mice were food restricted to 85-90% baseline body weight prior to and during testing to increase baseline responding. Mice were tethered to a 50 mm bilateral patch cable attached to an optical rotary joint (Doric), connected through an FC connector to laser (473nm, Shanghai Laser & Optics Century). Mice were placed in operant chambers (Med Associates) controlled by MedPC IV software. The beginning of each session turned on the house light, played a 0.5 s tone (2 kHz), and turned on LED cue lights located over each of the 2 nose ports. Each nose port contained photobeams and were baited with one 20 mg sucrose pellet (BioServ F0071) prior to each session. Beam breaks into each nose port (‘nose poke’) triggered a 0.5 s tone and turned off the LED cue lights for 1 s. Beam breaks into the active nose port also delivered laser stimulation (1 s, 10 mW, 40 Hz, 5 ms pulse width) through a TTL-generator controlled by an Arduino board. Nose pokes that occurred during the 1 s laser stimulation were recorded but had no consequence. Sessions lasted 45 min. Starting on day 5 of testing, the active nose port delivering laser stimulation was switched to the opposite nose port location (formerly the inactive nose port) until testing ended on day 8.

### Real-time place preference/avoidance procedure

Mice were tethered to a bilateral patch cable attached to an optical rotary joint (Doric) connected to a laser. During a baseline (no-laser) session, mice were placed on the border between two adjoining homogenous grey compartments (20 × 20 cm each). The amount of time spent in each compartment as well as entries into each compartment were recorded using AnyMaze software (v6; San Diego Instruments).

Most mice did not display a side preference, but any mice displaying a >75% side preference during the baseline session were excluded from further study (n=0). Starting on day 2, one side was designated active, wherein entries triggered laser stimulation (continuous, 473nm, 10mW, 40Hz, 5ms pulse width) controlled by AnyMaze software. Sessions lasted 20 min, and on days 5-7 the active (laser-delivering) compartment was switched.

### Instrumental sucrose task

Mice were placed into operant chambers (MedAssociates), equipped with two levers on either side of a magazine containing a food dish, and controlled by MedPC IV software. At the beginning of a session, house lights and LED cue lights above each lever were turned on. Lever presses at the active lever delivered a sucrose pellet (20 mg, BioServ F0071) into the food dish at a fixed ratio 1 and turned off the LED cue light for 1 s. Lever presses at the other, inactive lever, were recorded but had no consequence. Active lever location was the same across days for each mouse, but counterbalanced between mice. Sessions lasted 30 min.

### Open Field

Mice were placed into an open field (45 cm W x 45 cm D x 40 cm H) for 20-min sessions. Distance travelled, time in center, and number of entries into the center of the apparatus were collected using AnyMaze software.

### Statistical Analyses

Data were analyzed using Student’s t-test, 2-way repeated-measure ANOVA followed by Sidak post-hoc multiple comparisons, or Pearson correlation (GraphPad Prism v6). All data are represented as mean ± standard error of the mean (SEM) and/or as individual points.

## Acknowledgments

We thank Nick Hollon for guidance on analysis of fast scan cyclic voltammetry results. This work was supported by grants from NIH (R01DA036612, F32MH122192, P30DA048736), and Veterans Affairs (I01 BX003759).

## Figures

**Supplemental Figure 1.**
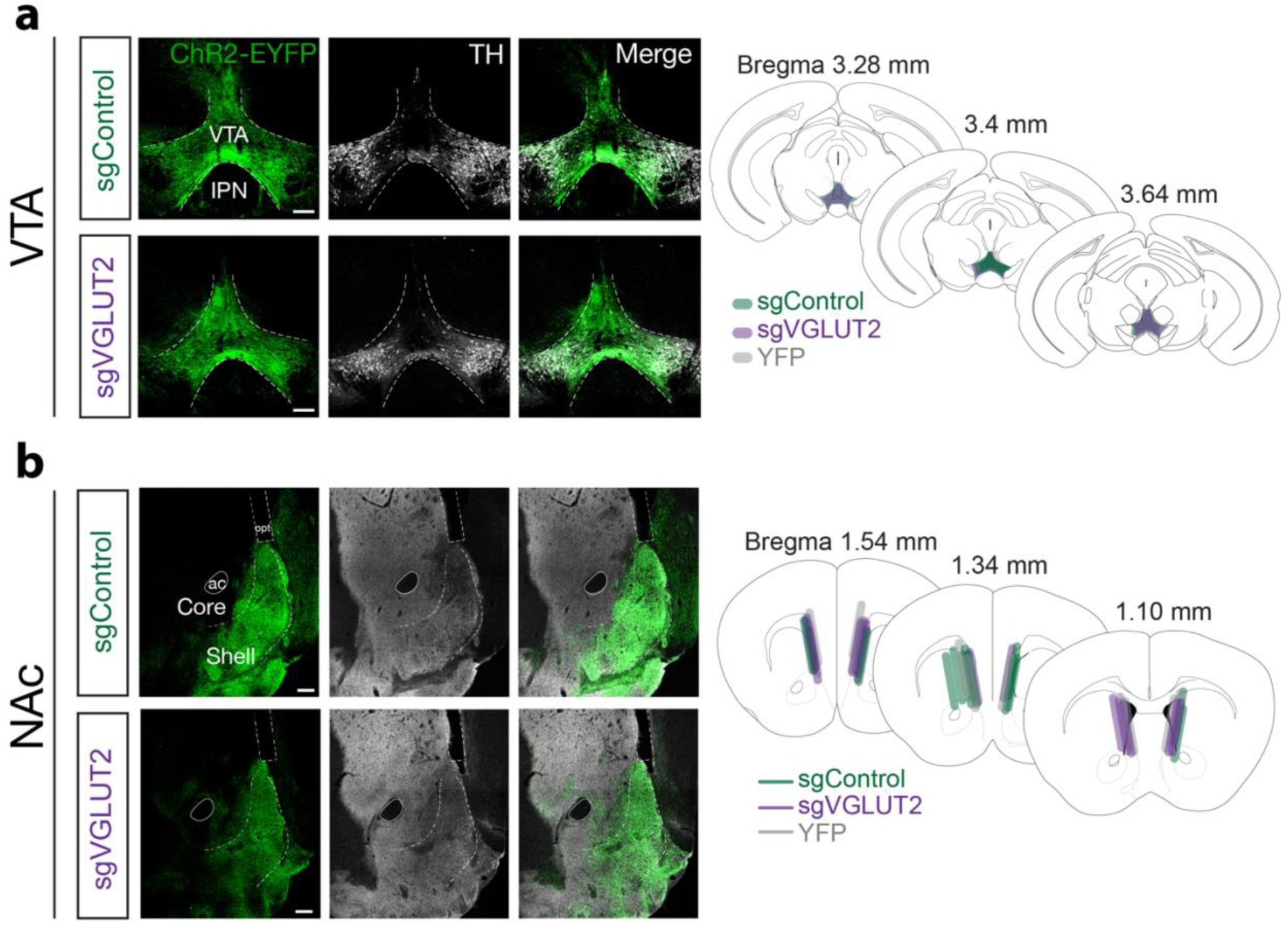
ChR2 expression and map of optic fiber placements. a) Images of coronal sections showing ChR2 (ChR2-EYFP, green) and TH (white) expression in VTA of sgControl and sgVGLUT2 mice. Scale bar=200 μm. Right shows ChR2 spread throughout VTA of each mouse. b) Images of coronal sections showing ChR2 (ChR2-EYFP, green) and TH (white) expression in NAc of sgControl and sgVGLUT2 mice. Scale bar=200 μm, anterior commissure (ac), optic fiber track (opt). Right shows bilateral placements of optic fibers in NAc of each mouse.

**Supplemental Figure 2.**
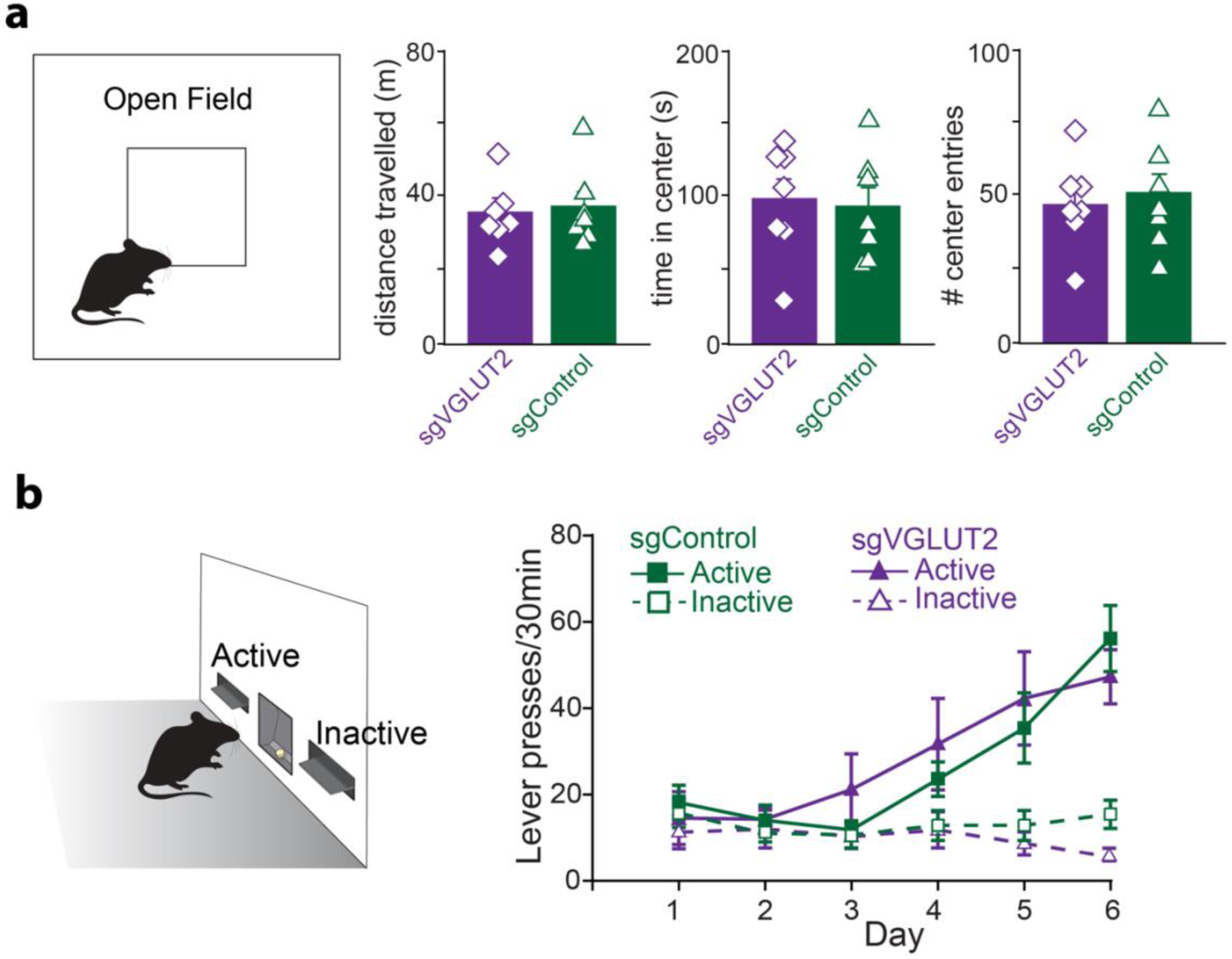
VGLUT2 deletion in VTA glutamate neurons does not affect baseline locomotion or reward learning. a) Schematic of open field test depicting center area (center square) and outside area. Distance traveled (unpaired t-test, t_12_=0.42, p=0.65), time in center (t_12_=0.15, p=0.75), and entries into center (t_12_=0.27, p=0.71) were all similar between sgVGLUT (n=7) and sgControl (n=7) mice. b) Schematic of instrumental sucrose self-administration assay. Presses at the active lever earned a 20 mg sucrose pellet (FR1 schedule) while inactive lever presses earned nothing. c) Active vs. Inactive lever presses for sgControl (n=7) and sgVGLUT2 (n=7) mice across daily training sessions showed both groups discriminated between active and inactive levers at similar levels (main effect of group: F_2,17_=0.81, p=0.46).

**Supplemental Figure 3.**
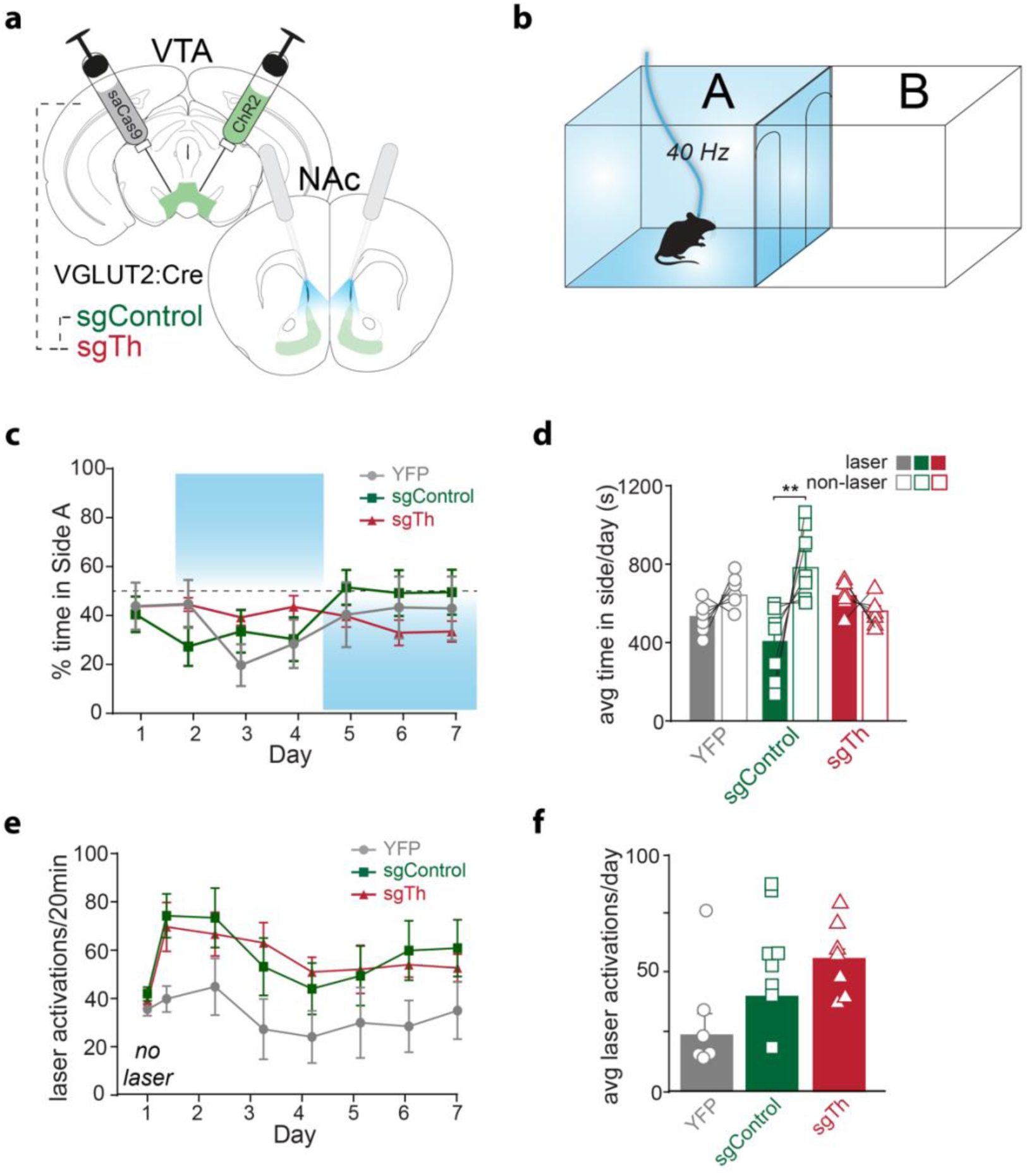
VTA glutamate projections require dopamine co-release to promote place avoidance. a) Illustration of injections in VTA and optic fiber placements above NAc. b) Schematic of RTPP assay. c) Percent time in Side A during each 20 min daily session. No laser was delivered on day 1, and location of laser side was reversed on day 5. d) Average time spent in laser side was lower than time in non-laser side among sgControl mice (n=8; Holm-Sidak’s multiple comparisons test, t=4.1, p<0.01) but not among sgTh mice (n=7; t=0.89, p>0.05), or YFP mice (n=6; t=1.3, p>0.05). e) Laser activations earned during each daily 20-min session. f) Laser activations averaged across laser sessions in sgTh mice (n=7), YFP mice (n=6) and sgControl mice (n=8).

